# Evaluation of a guideline developed to reduce HIV-related stigma and discrimination in healthcare settings and establishing consensus

**DOI:** 10.1101/333229

**Authors:** Garumma Tolu Feyissa, Craig Lockwood, Mirkuzie Woldie, Zachary Munn

**Author notes:** Corresponding Author E-mail address (GTF).

## Abstract

**Background:** Developing guidelines and policies is critical to address HIV-related stigma and discrimination (SAD) in healthcare settings. To this end, a multidisciplinary panel developed a guideline to reduce SAD. This project evaluated the appropriateness of implementing the guideline in the Ethiopian context.

**Methods:** A consensus of the expert panel was established through a Delphi technique which was followed by a panel meeting. Initial tentative recommendations were distributed to experts through e-mails to be evaluated using the modified guideline implementability appraisal (GLIA) v.2.0 checklist.

**Results:** In the first round of the Delphi survey, all (13) panel members evaluated the guideline. The overall score for the general domain of the modified GLIA checklist was 96.56%. The scores for individual recommendations ranged from 68.33% to 92.76%. Maximum and minimum scores were attained for measurability (97.71%) and flexibility (59.77%) domains respectively. Percentages mean score lower than 75% was obtained for flexibility and validity domains. Participants suggested that additional tools and training should be added to the guideline. In the second round of the survey, all the recommendations received endorsement with scores above 75%. Maximum and minimum scores were attained for measurability (100%) and flexibility (86.88%) domains respectively. During the panel meeting, issues of responsibility for implementing the guideline were discussed.

**Conclusion:** The project evaluated implementability of a guideline developed to reduce HIV-related SAD in healthcare settings. The Delphi survey was followed by a half-day meeting that helped in further clarification of points.

## Background

People living with the human immunodeficiency virus (HIV) are confronted with the physical, psychological and social impacts of the disease [1-5]. Stigma and discrimination (SAD), also called the “third phase of HIV/AIDS epidemics”, have been among the obstacles challenging actors working on the prevention and control of HIV [6]. SAD related to HIV are manifested in various forms such as: differential care or refusal to treat, testing and disclosure of the sero-status of clients without consent, verbal abuses or gossip, marking the files of patients, isolating them and excess use of precautions [7, 8].

The limited awareness of SAD, how they manifest and their consequences, prejudicial and stereotypical attitudes related to gender identity and sexual activity, and fear of HIV transmission are among factors contributing to SAD in healthcare facilities [9]. Hence, developing appropriate guidelines, policies, and redress systems and appropriate orientation of the rights and responsibilities of HCWs and patients are critical [9]. Cognizant of this, as described in the previous chapter, we have systematically developed a list of working recommendations to reduce SAD in healthcare facilities. Systematically developed guidelines are the source of summarized information [10]. Nevertheless, the development of guideline recommendations by itself is not enough. Other factors such as environmental and contextual factors need to be considered before making final decisions on the implementation of the guideline [11, 12].

Some researchers argue that using a theoretical framework will help to systematically identify and address factors that hinder guideline implementation [13, 14]. Factors such as reviewing, reporting and publishing guidelines have been found to enhance the implementation of the guidelines.[15] On the other hand, Jordan et al. argue that dissemination should involve an active process apart from the mere publication of guidelines [10]. Moreover, before officially publishing or disseminating a guideline, internal and external evaluation is required to promote the uptake the guideline [16, 17]. In addition to the development of tools to assess the rigor of the guideline development process [18], researchers have developed tools that help to assess both the rigor and implementability of guidelines [19]. Guideline developers and experts recommend assessing recommendations included in practice guidelines using guideline implementability checklists to make sure that the recommendations are clear and easy to implement [16, 17].

As described in chapter five, we have developed guideline recommendations based on an analysis of global evidence retrieved through literature searching. Therefore, this project aimed to assess the clarity, acceptability, implementability and relevance of the current guideline using Guideline Implementability Appraisal (GLIA version 2.0) checklist [20].

The objective of this project was to evaluate the appropriateness of the guideline developed to reduce HIV-related S&D to be implemented in the Ethiopian context. Specifically, the project aimed:

- To evaluate the appropriateness of the guideline to the Ethiopian context through a multi-round of Delphi surveys among the guideline panel.
- To evaluate the appropriateness of the guideline through a survey of external experts.
- To make amendments to each recommendation included in the guideline based on the comments of the experts.

## Methods

This project assessed the drafted recommendations for feasibility and appropriateness to the Ethiopian context. Consensus of the experts engaged in the evaluation was established through a modified Delphi technique [21].

### Rationale for the use of the Delphi technique in this project

The Delphi technique involves a series of questionnaires that are used to test opinion consensus amongst a group of experts [22, 23]. The technique can be conducted by email, online surveys or by post [23]. It is a preferable method of choice when there is little evidence regarding the topic, when participant anonymity is required, and when the cost and practicalities of bringing the participants together is prohibitive [24]. By assuring anonymity, it reduces the effect of dominant individuals and unwillingness to abandon publicly expressed opinions [25]. The Delphi technique also reduces reluctance to mention opinions that are unpopular, disagree with one’s associates, modify previously stated positions [26].

The choice for the specific type of consensus method is determined by the purpose of the study, the availability of scientific evidence in the field, the model of participant interaction, time and costs [24]. The aim of the current project was to translate research evidence into practice through the development of an evidence-informed guideline based on the consensus of experts. The development of a guideline needs a rigorous process to achieve consensus of experts. The Delphi technique is supposed to be more suitable compared to other consensus building methods [27]. This project sought the opinion of experts by keeping their responses anonymous and allowing them to freely express their opinions through e-mail surveys. In addition, the technique gave adequate time to the experts to exhaust options before making decisions. Hence, a modified Delphi technique was selected as a method of establishing consensus.

The Delphi technique is a hybrid of qualitative and quantitative methods [28]. The technique has been used in health disciplines since the 1970s [22]. It has been used by researchers to translate scientific knowledge and professional experience into informed judgment, in order to support effective decision-making [29]. The Delphi technique has been reported to be the most widely used consensus method for developing clinical guidelines [30-32]. Delphi techniques have been used to develop guidelines, to establish consensus on the use of the guidelines and to establish and evaluate how well a clinical practice is conforming to guidelines [33]. In the current project, the Delphi technique was used to establish consensus on the use of the each of the recommendations that constituted a guideline to reduce HIV-related stigma and discrimination in Ethiopian healthcare settings.

### Delphi process

The Delphi procedure starts with the selection of experts and is executed in a series of rounds [24, 25]. In a Delphi survey, appropriate selection of experts is essential for ensuring the quality of the data and increasing response rates. There is no standard definition of expert[27] and the definition depends on the specific objective of the research [27], but in general an expert is someone who has some knowledge of a specific subject [27, 34]. In the current project, experts were people who were knowledgeable of the subject matter by virtue of their role as clinicians with HIV patients, managers for HIV programs or researching on HIV. Experts may be selected based on records of relevant publications, their relationship with the topic and institutional positions they hold [35]. In the current project, the judgment of expertise was made based on their contribution in the field. Hence, researchers with relevant research projects and publications; health service managers and health professionals working on clinical or programmatic areas of HIV were selected as members of the guideline working group and experts for the current Delphi study. Apart from their expertise, the availability and commitments of the experts in the field were considered in selecting the panel members. The snow balling method was used to identify the experts. Finally, experts who were willing to participate were included in the multi-round survey.

### Panel size

There is no consensus on the panel size required for Delphi studies [29, 36]. Different Delphi studies have used different sample sizes ranging from as small as five to as large as 2865 [29]. In the Delphi technique, sample size does not depend on statistical calculations; rather it depends on the dynamics of arriving at consensus [37]. Some experts in Delphi techniques recommend careful selection of the panel for the specific topic of interest instead of increasing sample size or making the sampling process random [38]. In this project, 13 experts accepted my invitation and participated in the survey.

As a facilitator of the Delphi technique, the principla investigator set deadlines for each round of the Delphi and he used e-mail reminders for non-responders as an additional mechanism for increasing the response rate. The principal investigator sent the e-mail reminders three days after the deadline [35]. Respondents were given a three-week period for each round of Delphi [35]. As in other Delphi techniques, the opinion of every group member was reflected in the final group response [26]. The statistical average of the final opinions of the individual members was used to define group opinion [26].

### Data collection

After obtaining the list of experts, the principal investigator (GTF) made initial contacts to all experts giving them the purpose and procedures involved in the project and requesting them to participate in the development of the guideline. After receiving consent, the principal investigator sent the experts initial tentative recommendations by e-mail. Experts were asked to comment on each recommendation. We analyzed and summarized both qualitative and quantitative responses [24, 25].

There are three options to start Delphi round one. The first option is where Delphi round one is conducted as a qualitative study using open-ended questions to develop quantitative tools for the successive rounds [39]. In this approach, the first round is used to identify issues to be addressed in later rounds. The second option is where qualitative data can be collected through focus groups or interviews before the Delphi study and used to inform a quantitative first round of the Delphi [27]. The third option is where the quantitative first round is informed through a literature review or clinical practice [27, 40]. The first approach is often used in a classical (original) Delphi [41]. The second and third approaches are usually used in a modified Delphi technique [41].

In the current project, the tentative recommendations were informed by a systematic literature search and content analysis of the evidence. In this project, the modified Delphi, sometimes called ‘e-delphi’[41] was used. The purpose of the modified Delphi technique in this project was to get a consensus among the guideline panel on the tentative recommendations, and to modify the recommendations based on the responses of the experts. Therefore, the third approach was employed. Hence, experts were asked to rate each tentative recommendation using the Guideline Implementability Appraisal (GLIA V.2.0) checklist [20]. The GLIA checklist has options for both close-ended responses and open-ended responses. Hence, in addition to rating the recommendations, the panelists were asked to provide their suggestions on how to improve the implementations, feasibility and/or wordings of the specific recommendations. Participants were also encouraged to comment on the main guideline using track changes and highlights. The GLIA v.2.0 checklist was modified and used to assess the implementability of the guideline [20]. The GLIA v.2.0 [20] instrument contains 30 items in nine domains: global quality, executability, decidability, validity, flexibility, effect on process of care, measurability, novelty and computability [20]. Out of these, the last domain (computability) is used when there is a plan for electronic implementation [20]. Since this will not be part of the current work, the four items in this domain were not included in the questionnaire.

In this project, a modified GLIA v.2.0 checklist was used to assess the implementability of the guideline. The comments provided by the experts were incorporated into the successive round of the Delphi. In the subsequent round of the Delphi, we asked participants whether they would agree with the modified recommendations [24, 25]. We sent additional ideas in each round of the Delphi to the experts in the respective subsequent rounds [24, 25].

There is no template indicating the exact number of rounds needed for a Delphi study. Such decisions are pragmatically made by the researcher. Hence, the procedure is reiterated until the stability of responses is achieved [25]. Stability of responses is defined as “the consistency of responses between successive rounds of a study”[42](pp.84). Dajani et al. recommends measuring the level of agreement only if a stable answer is reached [42]. For each recommendation, once stability of the responses is achieved, consensus will be established [42].

For this project, recommendations having a general agreement of 75% and above were incorporated into the guideline. Recommendations with a rating lower than 75% were considered for modification to be incorporated into the subsequent rounds based on the comments of the respondents. In addition, specific comments given for each recommendation were considered for making modifications, adding or dropping a recommendation. The Delphi series stopped after stability was achieved (a 75% level of agreement) for each recommendation and if no newer comments emerged. The Delphi process is normally expected to achieve both consensus and non-consensus [43]. Therefore, in the current project, recommendations for which experts consistently disagreed were excluded or modified.

### Data quality control

In Delphi techniques, the opinion of every group member is reflected in the final group response [26]. Since decisions are made based on opinions of groups in the real world, Delphi techniques are believed to provide evidence of face validity [44]. In addition, Delphi is conducted in successive rounds, contributing to concurrent validity of the findings [45]. Researchers also believe that a Delphi technique provides reliable findings, because it achieves interaction among experts and at the same time avoids individual influences. Delphi overlaps both interpretive/qualitative and positivist/quantitative paradigms. Hence, researchers recommend the use of the term ‘trustworthiness’ to establish rigor in a Delphi study.[46] The concept ‘trustworthiness’ encompasses credibility, transferability, dependability and confirmability [47]. In Delphi studies, credibility is established by ongoing iteration and feedback given to the experts [48]. Therefore, the very beginning of the Delphi process makes it credible. In this project, dependability was enhanced by including relevant experts in the field.[49]. Confirmability is achieved through the collection of thick descriptive data, negative case analysis and arranging for a confirmability audit and establishing referential adequacy [47]. In this project, we kept accurate records of participants’ comments and responses in each round. We sent the comments of experts to the panelists in subsequent rounds. In addition, there was a face-to-face meeting prepared for further clarification. The transferability of an evidence is based on the similarity of contextual factors in the settings [50]. Therefore, other researchers and guideline implementers or developers were advised to take the consideration of the similarities of their respective contexts with the current situation and the current context of Jimma University Medical Centre (JUMC) when considering the potential transfer of the evidence into other settings.

### Data analyses

We conducted qualitative content analysis of the comments and we used the result of the analysis to modify the recommendations. In addition, we conducted the following quantitative analyses:

1. Percentage response rates,
2. Percentage scores for each domain of GLIA V.2.0: the total score for each GLIA domain was calculated by summing up total scores for all panel members. Then, the percentage score was obtained by dividing the total score by the maximum possible score.
3. Percentage agreement for each recommendation was calculated for each round of the Delphi. This information was used to modify recommendations, especially those with endorsement of less than 75%. In the cases where experts did not describe reasons for non-endorsement and for controversial issues, discussions on the recommendations were made through face-to-face meetings amongst the panel.

In this project, we wanted to take into consideration the input of each member of the panel. Instead of taking individual responses as outliers and rejecting them, a mechanism was in place in which they would clarify their opinions, which opens up for further comment by other members of the panel. Moreover, the panel consensus data were complemented with external panel review.

## Results

A formal consensus was sought from all the panel members using two rounds of panel surveys and an external panel review. This section describes results of these surveys.

### First round Delphi survey

In the first round of the Delphi survey, all (13) panel members evaluated the guideline. The overall score for the general domain of the GLIA version 2.0 score was 112 (% of maximum possible score=95.73%). Maximum score was achieved for the measurability domain (96.65%) and the minimum score was recorded for the flexibility domain (59.97%). A percentage mean score lower than 75% was obtained only for two domains: flexibility and validity domains (Table 1). The experts provided comments on how to improve or why modifications were needed for individual recommendations included in the guideline. The comments given were categorized into:

a. **General comments:** Comments that were provided for the entire guideline. These comments were suggestions for additional tools and training that should be part of the guideline
b. **Comments on specific recommendations:** Comments questioning the clarity and feasibility of implementing the recommendations.

**Table 1:** Guideline implementability (GLIA V.2.0) domain scores.

| GLIA Domain | Internal evaluation |  |  |  |  |  | External evaluation |  |  |
| --- | --- | --- | --- | --- | --- | --- | --- | --- | --- |
|  | Round 1 |  |  | Round 2 |  |  |  |  |  |
|  | Mean | SD | %age score | Mean | SD | %age score | Mean | SD | %age score |
| Executability | 21.81 | 3.35 | 83.88 | 15.38 | 0.77 | 96.13 | 10.50 | 1.51 | 87.50 |
| Decidability | 33.1 | 3.36 | 84.89 | 23.85 | 0.38 | 99.38 | 17.25 | 1.54 | 95.83 |
| Validity | 17.48 | 4.77 | 67.23 | 15.85 | 0.554 | 99.06 | 10.50 | 1.93 | 87.5 |
| Flexibility | 23.39 | 4.23 | 59.97 | 20.85 | 0.38 | 86.88 | 9.66 | 1.37 | 53.70 |
| Effect on process of care | 24.71 | 1.04 | 95.04 | 15.77 | 0.44 | 98.56 | 11.33 | 0.89 | 94.44 |
| Measurability | 25.13 | 0.96 | 96.65 | 16.00 | 0.00 | 100 | 10.17 | 0.39 | 84.72 |
| Novelty | 33.44 | 2.14 | 85.74 | 23.08 | 0.49 | 96.08 | 17.00 | 1.41 | 94.44 |

NB: GLIA: Guideline Implementability Appraisal, SD: standard deviation, %age score: percentage score

The scores for individual recommendations ranged from 151 (68.33%) to 205(92.76). Six recommendations received an endorsement of lower than 75%. The recommendations with endorsement lower than 75% were:

1. **Counselling and behaviour change programs to address self-stigma (endorsement score=71.04%)** The most important reasons for the low score for this recommendation was described as lack of detailed description of the recommendations and failure to specify the type of behavioural change programs.
2. **Group intervention through telephone support for people living with HIV (endorsement score=68.33%)** The feasibility of this intervention was questioned by the panel. Therefore, this recommendation was brought for panel discussion during the second round panel meeting.
3. **Micro-finance and livelihood programs to create economic opportunities (endorsement score=70.14%)** Concern was raised because participants claimed that it was not the mandate of healthcare institutions to provide microfinance interventions and resource-wise, this recommendation was reported to be not feasible. Therefore, this recommendation was brought for panel discussion during the second round panel meeting.
4. **Training programs to gain facilitation skills, processes to collect and analyse data for advocacy (endorsement score = 70.14%)** This recommendation was rated a low score because of limited description linked with it. The feasibility of the recommendation was also questioned.
5. **Developing stigma and discrimination reduction policies with employees (endorsement score = 73.76%)** The panel requested description of this recommendation, specifically by linking with previous research findings.
6. **Programs, offices and institutions need to advocate temporary special measures such as affirmative action for women and special forums for participation (endorsement score=71.04%)**

The feasibility of this recommendation was questioned as it was perceived by some panel members to be beyond the scope of health institutions. In addition to the above comments targeting individual recommendations, as mentioned in table 2, the panel suggested that some recommendations should be merged. The main comments made by the panel during the first round survey are summarized in Table 2. Based on the first round comments, modifications were made. The second round survey was then conducted after incorporating comments from the first round and modifications to the guideline.

**Table 2:** Summary of comments provided during first round survey.

| S/n | Comments | Actions/resolution |
| --- | --- | --- |
| **General comments** |  |  |
| 1. | The sequence of applying these recommendations is not clearly documented | This has been indicated at the end of the recommendations incorporating steps in implementation |
| 2. | In the introduction part, the intended audience should also include non-health disciplines such as psychology and sociology who work to improve the psychosocial well-being of PLHIV | Accepted |
| 3. | For most of the recommendations: patient characteristics (co-morbidities) were not mentioned | Most recommendations work for all types of HIV patients regardless of their co-morbidities |
| 4. | Settings such as faith-based organizations may be included as part of the guideline | This is beyond the scope of the current guideline, which is limited to healthcare settings |
| 5. | The guideline should be broad, and the scope should be beyond the health sector | This cannot be addressed within the time frame. After this project is over, we may consider developing guidelines for other settings |
| 6. | Additional tools should be part of the guideline | Accepted and added tools to be posted and tools for monitoring and evaluation |
| 7. | Key population should be defined | Accepted |
| 8. | People associated with the virus should be defined | Accepted |
| 9. | Stigma occurs when those health care workers who are not aware of HIV-related stigma provide services to HIV patients. Therefore, the type and role of service providers needs to be specified. | Brought for discussion by the panel during the second meeting and further explored during key informant interviews |
| **Comments on specific recommendations** |  |  |
| 10. | RN1.4, RN2.4 and RN3.4 are fragmented and can be better strengthened if they are merged together. | Accepted and merged the recommendations<br>So, RN1.4, RN2.4 and RN3.4 were merged |
| 11. | RN2.1 should be supported with evidence | Accepted, reference and quality of evidence included |
| 12. | RN2.1 and RN2.2 can be merged | Accepted |
| 13. | RN 2.2. is not detailed | Accepted |
| 14. | RN2.3 is not detailed. Group support through telephone is not clear enough. Are you going to call them or text them through SMS? It is not feasible, and the quality of evidence is also very low. Also, it is better to use references | Brought for discussion by the panel during the second meeting |
| 15. | One of the recommendations, micro-finance interventions is not feasible | Brought for discussion by the panel during the second meeting |
| 16. | RN2.4 needs resources | Suggestions will be sought from panel members on whether the allocation of such resources is feasible will be discussed |
| 17. | RN2.6 is not specific | The recommendation was dropped |
| 18. | RN2.6 is difficult to measure unless we put measurement parameters | Accepted |
| 19. | RN3.1 is not detailed | Accepted |
| 20. | RN4.2 is not detailed | Accepted |
| 21. | RN6.2 Needs details | Accepted |
| 22. | RN6.3 needs details | Accepted |
| 23. | RN1.4 is not feasible | Accepted |
| 24. | RN4.1. is not feasible | Accepted |
| 25. | Co-morbid mental illness among HIV clients plays critical role in worsening the stigma towards HIV patient. So, consider mental illness | This will broaden our scope. We may consider another guideline for this. |
| 26. | All recommendation need at least orientation and training | The guideline will be introduced through different methods including orientation and training. And additional methods will be further sought from the panel |
| 27. | RN1.2, RN 6.3 and 6.11 need to be merged or be described using a single recommendation. | Accepted |

**NB:** HIV: Human immunodeficiency virus, PLHIV: People Living with HIV, RN: Recommendation number

### Second round Delphi survey

Eight of the 13 (61.5%) panel members responded to the second round survey using the GLIA V.2.0 checklist. Five panel members did not provide ratings during the second round Delphi survey. Of these, four of them participated in the second round panel meeting. In the second round panel meeting, all the comments in the first round and second round were summarized and discussed. Hence, those members who missed the second round survey got the opportunity to reflect on their ideas in the meeting. In the second round, the general domain received an endorsement score of 64/72 (88.89%). Maximum score was attained for the measurability domain (100%). A minimum score was recorded for the flexibility domain (86.88%) (Table 1). In the second round of the Delphi survey, each recommendation received an endorsement of over 75%. Only a few comments were raised by the panel. The summary of the comments and the respective resolutions made following the comments is shown in Table 3.

**Table 3:**
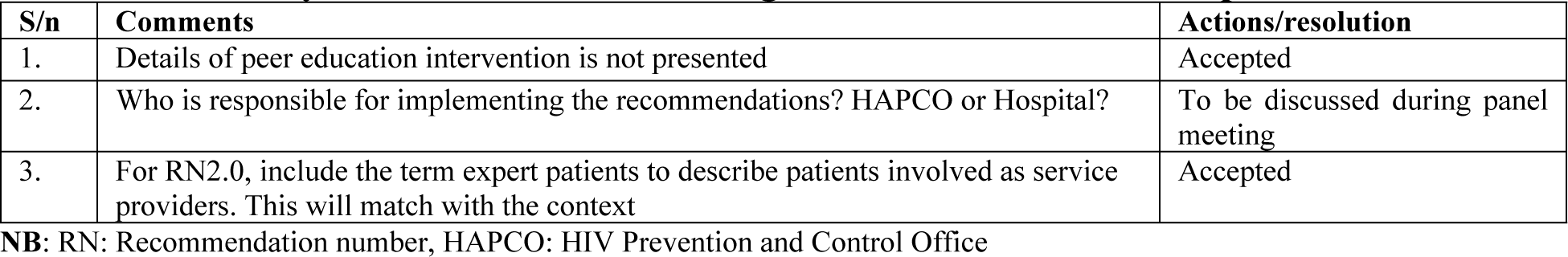
Summary of comments made during second round and the respective resolutions.

| S/n | Comments | Actions/resolution |
| --- | --- | --- |
| 1. | Details of peer education intervention is not presented | Accepted |
| 2. | Who is responsible for implementing the recommendations? HAPCO or Hospital? | To be discussed during panel meeting |
| 3. | For RN2.0, include the term expert patients to describe patients involved as service providers. This will match with the context | Accepted |

NB: RN: Recommendation number, HAPCO: HIV Prevention and Control Office

The highest percentage mean score was attained for the measurability domain (100%) and the lowest mean score percentage was attained for the flexibility domain (88.88%) (Table 1). All individual recommendations received endorsements with scores over 75%. Since there were few comments given in the second round survey and the ratings for the recommendations were also high, the panel decided not to have additional surveys. Instead, a second round panel meeting was called to discuss in person the comments made thus far and the modifications made. Further comments were sought from the panel. Major points raised during the meeting are briefly presented below.

### Major points of discussion during the second round guideline panel meeting

1. **The responsible body for implementation of the interventions should be clearly specified**: Based on detailed discussions, the panel resolved that all health professionals, healthcare facility administration and HIV prevention and control offices are responsible for the interventions in the recommendations be included in the guidelines. The panel recommended that training should be provided for those PLHIV who provide psychosocial support, adherence support and peer support for PLHIV.
2. **Whether microfinance intervention can still be part of the guideline:** The panel decided that HAPCO and healthcare facilities can routinely link patients to support organizations. Nevertheless, they agreed that it is very difficult for them to provide financial interventions, such as microfinance interventions.
3. **Whether telephone support interventions are still feasible for the context:** The panel resolved with the consensus that in the Ethiopian context, there is no adequate evidence indicating that such interventions are feasible. However, they all agreed that these interventions (phone calls and reminder texts) can be included as alternative methods for the provision of psychosocial support.
4. **Who is responsible for informing the rights and responsibilities to patients?** The panel resolved with the consensus that all health professionals should routinely inform patients about the details of procedures, their rights and responsibilities. In addition, healthcare facility administration and HIV Prevention and Control Office (HAPCO) are responsible to make sure that information is provided to patients on their rights and responsibilities. This information should include the rights that each patient has regardless of his or her sex, disease status, age and other characteristics.
5. **Whether translating the guideline into local language is needed:** The panel decided that for healthcare professionals, there is no need to translate the guideline into local languages. Nevertheless, the training manual that may be prepared in the future for peer supporters and expert patients (non-professionals) should be translated into local languages.
6. **Arrangement of recommendations** The panel suggested that the recommendations should be arranged, not under guiding principles, but under major thematic areas as conceptualized in the systematic reviews presented in previous chapters.

### Evaluation by external experts

Of the 13 experts invited to participate in the evaluation, six agreed to evaluate the guideline using the same checklist that internal evaluators used. The external experts gave an overall score of 51 (94.44%) to the general domain of GLIA. Each recommendation received an endorsement over 75%. The maximum score was recorded for the decidability domain (95.83%) and minimum score was attained for the flexibility domain (53.70%). The external panels did not provide many comments. Major comments made were categorized under general comments, comments specific to individual recommendations and comments related to format of the guideline (Table 4).

**Table 4:** Summarised comments from the external panel.

| S/N | Comments | Resolution/actions |
| --- | --- | --- |
| **General comments** |  |  |
| 1. | Recommendations should be action-oriented rather than descriptive. Some recommendations are not identifiable because of long descriptions | Accepted |
| 2. | Settings in which the guideline is to be implemented is not clearly described | Accepted |
| 3. | The guideline mainly focuses on the provider or user of the guideline and simply highlights the target. The targets must be described in detail in a separate section. | Accepted |
| 4. | Target organizations for the guideline are not mentioned except on the cover page. | Accepted |
| 5. | The required service modifications are not mentioned | Accepted |
| **Comments related to the format of the guideline** |  |  |
| 6. | Boxes for strategies and recommendations need to be separate | Accepted |
| 7. | Indicators need to be presented clearly for recommendations | Accepted |
| 8. | It is better to put boxes and tables at the end of description rather than putting them in the middle of text descriptions | Accepted |
| **Comments on specific recommendations** |  |  |
| 9. | RN3.3 does not show how opinion leaders execute their jobs | Accepted |
| 10. | RN43 does not detail how to empower PLHIV | Accepted |
| 11. | For RN61, RN62, RN63 AND RN64, strategies for implementation was not addressed well | Accepted |
| 12. | RN33 does not show logical sequences | Accepted |
| 13. | RN 51 is not detailed | Accepted |
| 14. | No evidence presented for RN61, RN62 | Accepted |

**NB:** PLHIV: People Living with HIV, RN: Recommendation number

## Discussion

This project attempted to evaluate a guideline to reduce HIV-related stigma and discrimination developed in chapter five, using guideline implementability appraisal (GLIA) version 2.0 checklist. The internal evaluation was conducted using two rounds of the Delphi survey that was followed by a face-to-face meeting of the guideline panel. The Delphi surveys were complemented by an additional evaluation by external experts.

In the first round Delphi survey, a percentage mean score lower than 75% was obtained for two domains: flexibility and validity domains of GLIA V2.0 checklist. This indicated that more work was needed with including detailed descriptions on areas such as strength and quality of recommendations and detailed justifications of recommendations. Therefore, modifications were made before sending the guideline for the second round evaluation. The modifications made were: incorporating strength and quality of recommendations for those recommendations for which such data were available.

As the experts involved in the Delphi survey were also members of the guideline working group, it was my expectation that the risk of dropping out from the study would be minimal. Nevertheless, in the second round, we obtained a response rate of 61.5%, which was lower than my expectation. This is, however, an expected limitation of Delphi techniques [22]. In addition, it is a common obstacle that guideline developers face when using the GLIA checklists as it is a long instrument and may result in low response rates [51]. However, the instrument provides an opportunity for a comprehensive evaluation of guideline recommendations. It helps to assess both implementability and rigor of recommendations [51].

The other potential reason for delayed responses and low response rates in the current project might be because the experts were occupied with other tasks and that the current project was conducted within a tight schedule. We had made efforts to reduce delays and drop outs by setting deadlines, e-mail and telephone reminders. Such mechanisms have also been used by previous researchers employing Delphi techniques [35]. On the other hand, the same experts who failed to provide responses for the second round survey participated in a panel meeting where they got an opportunity to reflect on their opinions. In the panel meeting, a summary of the comments and modifications made in all rounds were presented and reflections were made by all participants. Hence, the attrition bias related to drop outs was minimal.

During the external panel survey, the lowest score was recorded for the flexibility domain (53.70%). This was an indication that notified me to make the emphasis on the quality and strength of recommendations. This was a partially expected response as some recommendations still lacked quality and strength of evidence supporting them. Hence, for such recommendations, I indicated them as ‘no quality of evidence assigned’. Later, some of such recommendations were assigned as good practice points.

In addition, there was a concern by external reviewers regarding feasibility issues. Some enquired about the commitment of Jimma University Medical Center (JUMC) for availing continuous supply of materials for standard precautions. Therefore, this was later explored in detail during the key informant interviews (this is reported in chapter seven as part of contextualizing the guideline). On the other hand, the response ‘not applicable (NA)’ for question 18 might have contributed to the low score in the flexibility domain. The question enquires whether the recommendations were made with the consideration of co-morbidities among clients, which was not practical for the current guideline.

In general, except assigning a low endorsement score for the flexibility domain, the external panel endorsed all individual recommendations with scores above 75%. For the current Delphi survey, since some comments were merged, and additional new recommendations were added and dropped iteratively, it was not practical to employ statistical techniques such as weighted kappa, index of predicted association and McNemar chi-square tests [42, 52-54]. Nevertheless, additional comments were not forthcoming during the second round and during external expert evaluations. In addition, during the second panel meeting, a detailed discussion was held both on the comments and the modifications made to address the comments.

The current project employed a modified Delphi technique to establish consensus on recommendations based on the best available evidence from systematic reviews. Such techniques have been used by previous researchers to develop guidelines [55, 56]. One of the potential limitations of a modified Delphi approach is the absence of face-to-face engagement with panel members [55]. In the current project, this limitation was minimized by incorporating two panel meeting sessions, one before the start of the Delphi survey and one after the second round Delphi survey. This has helped to clarify and discuss vague points. However, before implementing the guideline, it is critical to identify contextual and environmental factors to tailor the implementation of the guideline to local context. In the next chapter (chapter seven) we have indicated how we explored details of contextual factors that potentially influence the implementation of the guideline.

## Conclusion

The current project evaluated the implementability of a guideline developed to reduce HIV-related stigma and discrimination in healthcare settings. The project employed both internal and external evaluation. The Delphi survey was followed by a face-to-face meeting that helped in further clarifications of points and addressing some of the limitations of the series of the Delphi surveys.

## Acronyms/Abbreviations

AIDS: Acquired Immune-Deficiency Syndrome
CDC: Centers for Disease Control and Prevention,
GLIA: Guideline Implementability Appraisal
HAPCO: HIV Prevention and Control Office
HIV: Human Immunodeficiency Virus
IRB: Institutional Review Board
JIH: Jimma University Institute of Health
JUMC: Jimma University Medical Centre
MDT: Multidisciplinary team
ORECI: Office of Research Ethics, Compliance and Integrity
PLHIV: People Living with HIV
SAD: Stigma and discrimination

## Declarations

### Acknowledgments

The authors are grateful to Siang Tay for editing the report.

### Availability of data and materials

All data were reported in the main manuscript.

### Authors’ contributions

Conceptualization: GTF, CL, MW and ZM.

Data curation: GTF, CL, MW and ZM.

Formal analysis: GTF.

Methodology: GTF, CL, MW and ZM.

Project administration: GTF.

Validation: GTF, CL, MW and ZM.

Visualization: GTF, CL, MW and ZM.

Writing – original draft: GTF.

Writing – review & editing: GTF, CL, MW and ZM.

### Ethical approval and consent to participate

The project has ethical approval both from the Institutional Review Board (IRB) of Jimma Institute of Health (JIH) at Jimma University (RPGC/389/2016) and the University of Adelaide Office of Research Ethics, Compliance and Integrity (ORECI) (approval number H-2016-140). Prior to the data collection, the objective of the research, potential harms and benefits of participating in the project were described to participants. Participants were provided with complaints procedure and information sheets, based on which informed consent was obtained. Anonymity of responses was assured by not disclosing the identity of participants.

### Consent for publication

Not applicable.

